## Supplementary Information for "α-catenin switches between a slip and an asymmetric catch bond with F-actin to cooperatively regulate cell junction fluidity"

#### Supplementary figures

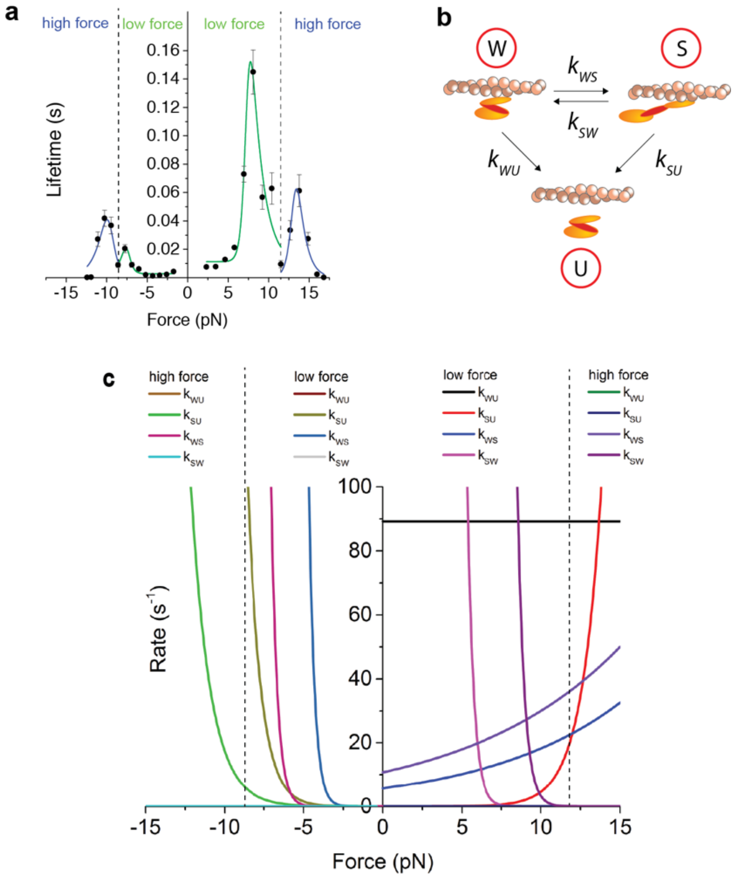


Supplementary figure 1: **Two-step catch-bond model**. **a**, The plot of the interaction lifetime between an α-catenin homodimer and actin vs force was divided into regions of low (green) and high (blue) force to separate the lifetime peaks. **b**, Within each force region, α-catenin reversibly transitions between a weak, folded bound state (W) and a strong, unfolded state (S), with force-dependent rates $k_{WS}$ and $k_{SW}$. Detachment from the bound states W and S to the unbound state U is described by the two rates $k_{WU}$ and $k_{SU}$, respectively. Force-dependent rates have the form $k_{ij} = k_{ij}^{0}\exp\left(\frac{d_{ij}F}{k_{B}T}\right)$, where $d_{ij}$ are the distance parameters and $k^{0}_{ij}$ are the rates at no load. **c**, Plot showing the force-dependent rates obtained from fitting the two-state model to data in panel (a). Numerical values of the fit parameters are reported in Supplementary Table 1a.

**a, single α-catenin homodimer**

F > 0 low force

| | Value ($s^{-1}$) | Standard Error ($s^{-1}$) | | Value (nm) | Standard Error (nm) |
|---|---|---|---|---|---|
| $k^0_{WU}$ | 89 | 7 | $d_{WU}$ | 0.00 | 0.04 |
| $k^0_{SU}$ | 0.0014 | 0.00012 | $d_{SU}$ | 3.31 | 0.04 |
| $k^0_{WS}$ | 5.7 | 0.5 | $d_{WS}$ | 0.47 | 0.04 |
| $k^0_{SW}$ | 5.9E7 | 1.4E7 | $d_{SW}$ | -10.1 | 0.1 |

F > 0 high force

| | Value ($s^{-1}$) | Standard Error ($s^{-1}$) | | Value (nm) | Standard Error (nm) |
|---|---|---|---|---|---|
| $k^0_{WU}$ | 954 | 46 | $d_{WU}$ | 1.61 | 0.01 |
| $k^0_{SU}$ | 7.1E-8 | 0.5E-8 | $d_{SU}$ | 3.15 | 0.02 |
| $k^0_{WS}$ | 10.8 | 0.5 | $d_{WS}$ | 0.42 | 0.01 |
| $k^0_{SW}$ | 3.4E10 | 0.3E10 | $d_{SW}$ | -9.38 | 0.03 |

F < 0 low force

| | Value ($s^{-1}$) | Standard Error ($s^{-1}$) | | Value (nm) | Standard Error (nm) |
|---|---|---|---|---|---|
| $k^0_{WU}$ | 1000 | 1400 | $d_{WU}$ | 0.0 | 0.7 |
| $k^0_{SU}$ | 9.3E-4 | 4.5E-4 | $d_{SU}$ | -4.19 | 0.06 |
| $k^0_{WS}$ | 1.5E-5 | 2.8E-5 | $d_{WS}$ | -11.0 | 0.2 |
| $k^0_{SW}$ | 18000 | 30000 | $d_{SW}$ | 0.0 | 0.2 |

F < 0 high force

| | Value ($s^{-1}$) | Standard Error ($s^{-1}$) | | Value (nm) | Standard Error (nm) |
|---|---|---|---|---|---|
| $k^0_{WU}$ | 80000 | 15000 | $d_{WU}$ | 0.0 | 0.1 |
| $k^0_{SU}$ | 0.00477 | 0.00036 | $d_{SU}$ | -3.35 | 0.03 |
| $k^0_{WS}$ | 1.28E-6 | 0.25E-6 | $d_{WS}$ | -10.52 | 0.08 |
| $k^0_{SW}$ | 1.4E8 | 3.9E8 | $d_{SW}$ | 8.6 | 1.4 |

**b,multiple α–β-catenin homodimers**

F > 0 low force

| | Value ($s^{-1}$) | Standard Error ($s^{-1}$) | | Value (nm) | Standard Error (nm) |
|---|---|---|---|---|---|
| $k^0_{WU}$ | 110 | 20 | $d_{WU}$ | 0.0 | 0.1 |
| $k^0_{SU}$ | 0.061 | 0.008 | $d_{SU}$ | 3.27 | 0.07 |
| $k^0_{WS}$ | 0.52 | 0.15 | $d_{WS}$ | 6.0 | 0.2 |
| $k^0_{SW}$ | 1.3E6 | 0.4E6 | $d_{SW}$ | -5.0 | 0.2 |

Supplementary Table 1: **Two-step catch-bond model – fit parameters**. Force-dependent rates have the form $k_{ij} = k^0_{ij}\exp\left(\frac{d_{ij}F}{k_BT}\right)$, where $d_{ij}$ are the distance parameters and $k^0_{ij}$ are the rates at zero load. **a**, Parameters of the force-dependent rates obtained from fitting the two-state model to the lifetime of a single α-catenin homodimer interaction (Fig. 1c). **b**, Parameters of the force-dependent rates obtained from fitting the two-state model to the lifetime of multiple single α–β-catenin heterodimers interaction (Supplementary figure 6).

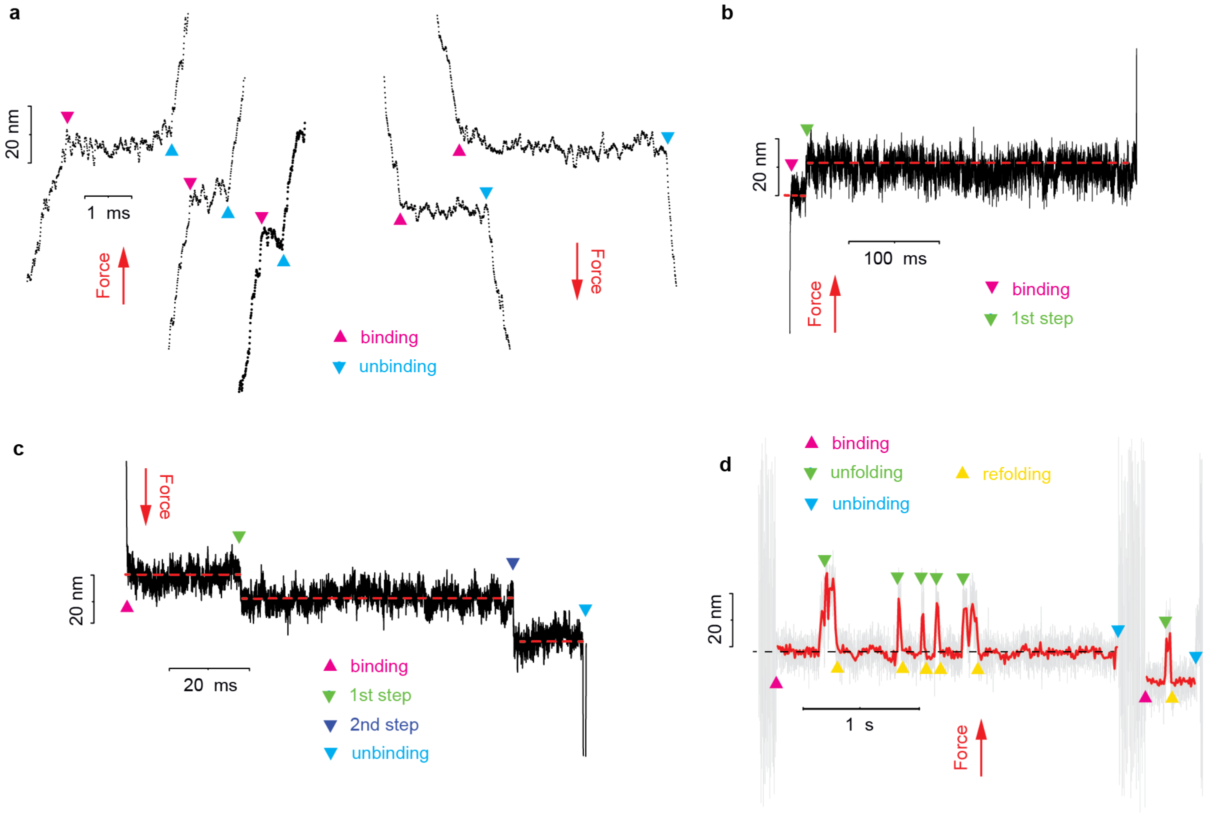


Supplementary figure 2: **Position records from α-catenin homodimers and α-β-catenin heterodimers under different forces**. **a,** An α-catenin homodimer under low force (~ 3 pN) shows single brief interactions; **b**, under moderate force (5 pN < F < 10 pN), α-catenin homodimer interactions show prevalently single steps; **c**, under high force (> 10 pN) α-catenin homodimer interactions usually display multiple steps. **d**, At about 5 pN force, a single α-catenin homodimer shows jumps between two positions.

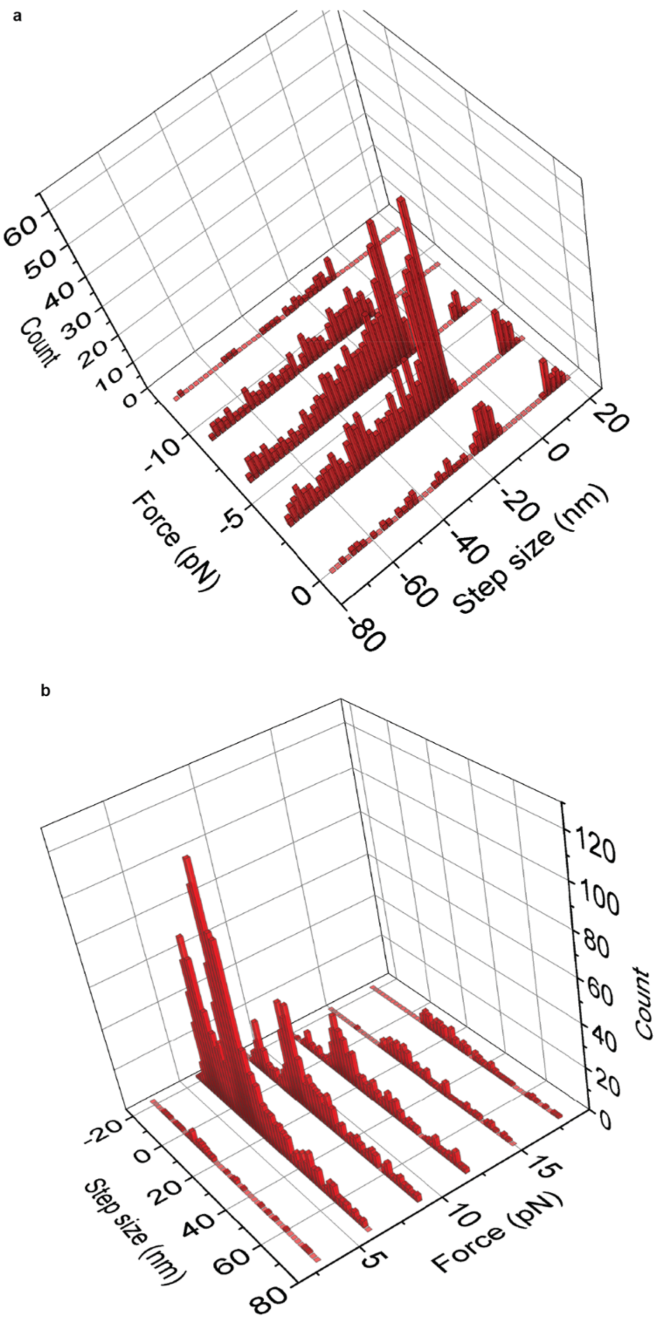


Supplementary figure 3: **Step size distribution versus force of a single α-catenin homodimer**. 3D plot of the step size distribution versus force for a single α-catenin homodimer molecule during its interaction with actin. **a**, negative forces. **b**, positive forces. The force sign is defined as in Figure 1.

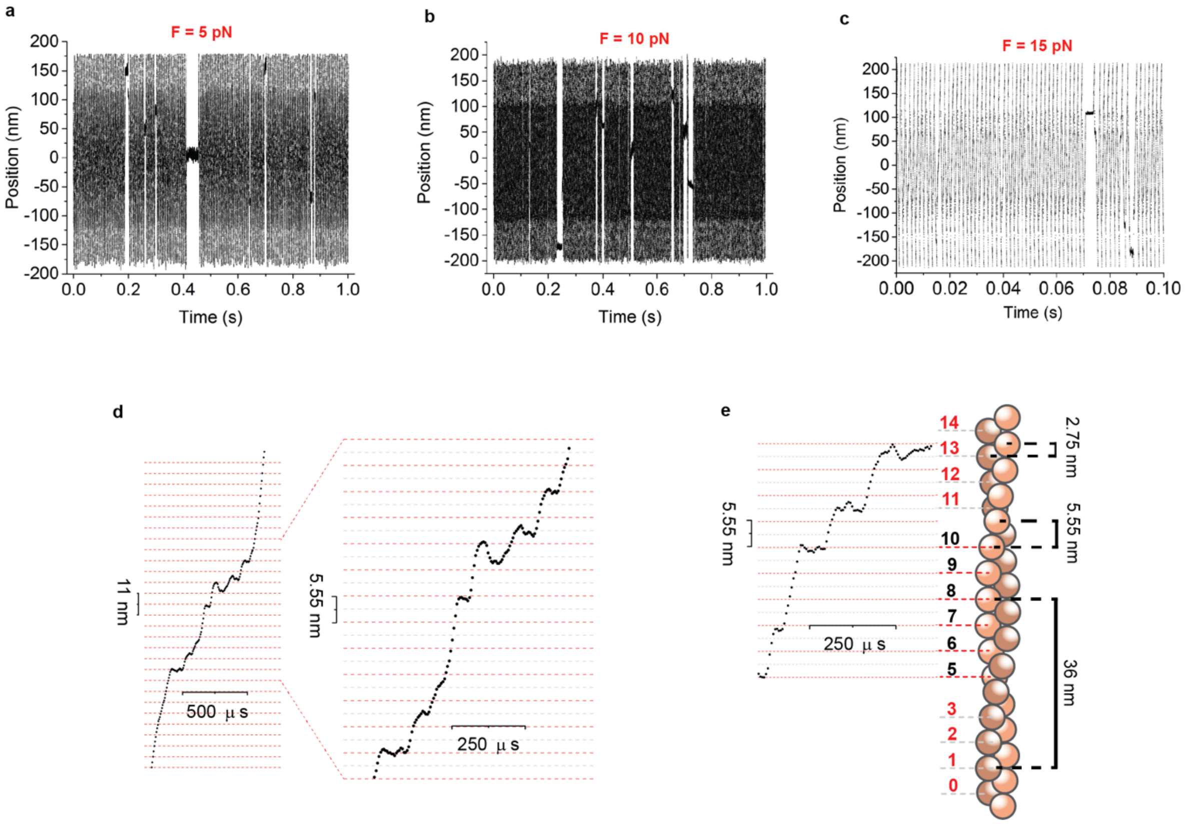


Supplementary figure 4: **Position records from single α-β-catenin heterodimers**. A single α-β-catenin heterodimer rapidly unbinds and rebinds from actin at all forces. Interaction lifetime decreases with force. **(a)** *F* = 5 pN, 1 s record; **(b)** *F* = 10 pN, 1 s record; **(c)** *F* = 15 pN, 0.1 s record. **(d)** Interactions are usually composed by a series of very rapid binding-unbinding sequences. As highlighted in Fig. 2d-f, binding occurs with a periodicity of 5.55 nm, which occasional shifts of 2.75 nm. **(e)** The figure shows that the binding periodicity of 5.55 nm is related to the distance between consecutive actin monomers, whereas the periodicity shift of 2.75 nm is a consequence of the distance between monomers on adjacent protofilaments. Traces in (d) and (e) are from experiments with *F* =15 pN.

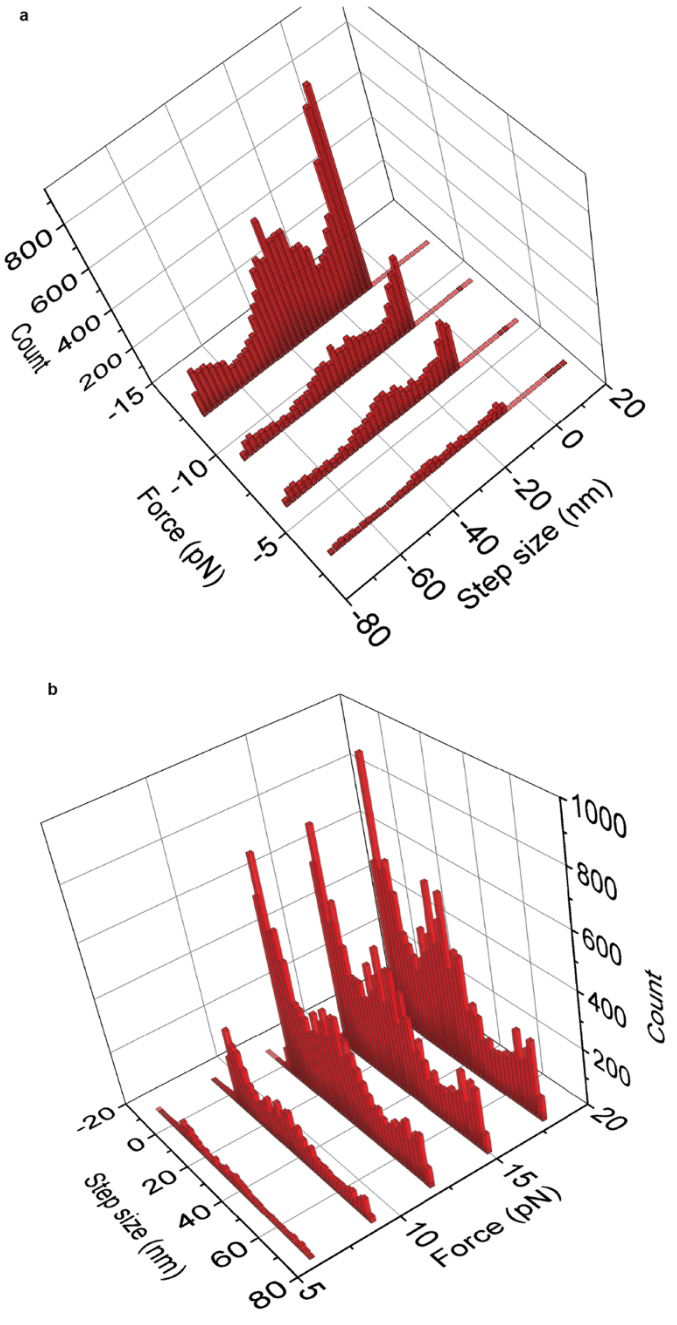


Supplementary figure 5: **Step size distribution versus force of a single α–β-catenin heterodimer**. 3D plot of the step size distribution versus force for a single α–β-catenin heterodimer during its interaction with actin. **a**, negative forces. **b**, positive forces. The force sign is defined as in Figure 2.

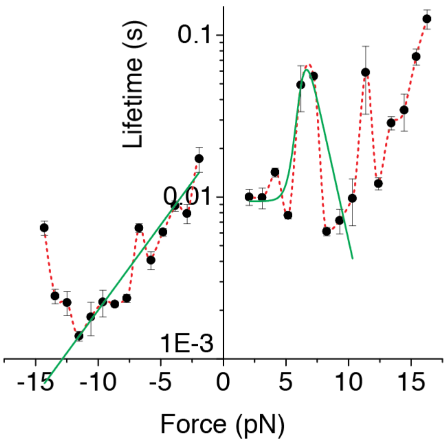


Supplementary figure 6: **Lifetime vs force for multiple α-β-catenin heterodimers**. Log-linear plot of the load-dependent lifetime of the interaction between multiple α-β-catenin heterodimers and actin. Green line is the fit of the peak occurring at lower force with the two-state catch-bond model at positive forces (see methods and Supplementary figure 1). Fit parameters are reported in the Supplementary table 1b. Fit of the peaks occurring at higher forces did not converge because of the low sampling of data. Blue line is the fit of lifetime τ with the Bell-bond equation $\tau = \tau_0 \exp\left(-\frac{d_{\alpha\beta}F}{k_B T}\right)$ (see Methods). Fitting parameters are $d_{\alpha\beta}$ = 1.0 ± 0.2 nm, $\tau_0$ = 23 ± 5 ms.

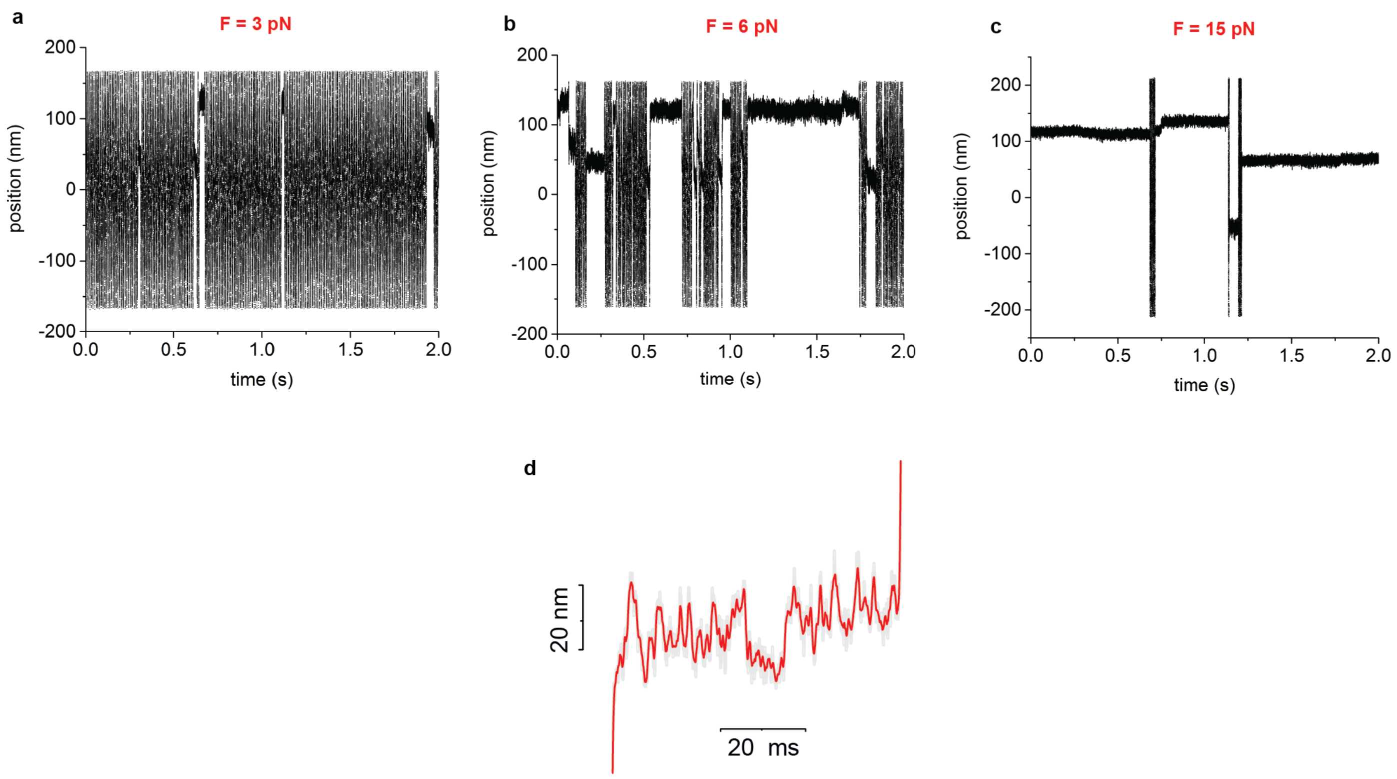


Supplementary figure 7: **Position records from multiple α-β-catenin heterodimers under different forces**. **a,b,c** Interactions of multiple α-β-catenin heterodimers with actin around 3 pN, 6 pN, and 15 pN, respectively. The lifetime of the interactions was significantly longer at 6 pN and 15 pN compared to 3 pN. **d**, Multiple α-β-catenin heterodimers at > 5 pN typically showed jumps between two positions.

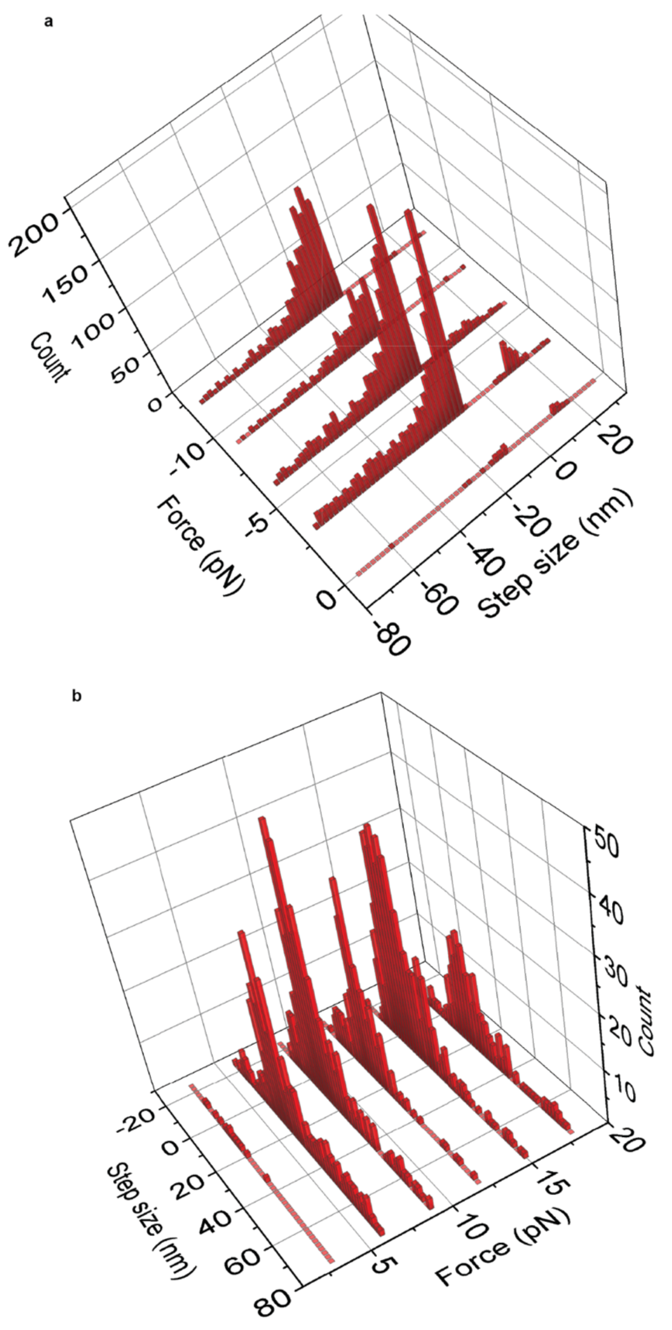


Supplementary figure 8: **Step size distribution versus force of multiple α–β-catenin heterodimer**s. 3D plot of the step size distribution versus force for multiple α–β-catenin complexes during their interaction with actin. **a**, negative forces. **b**, positive forces. The force sign is defined as in Figure 3.

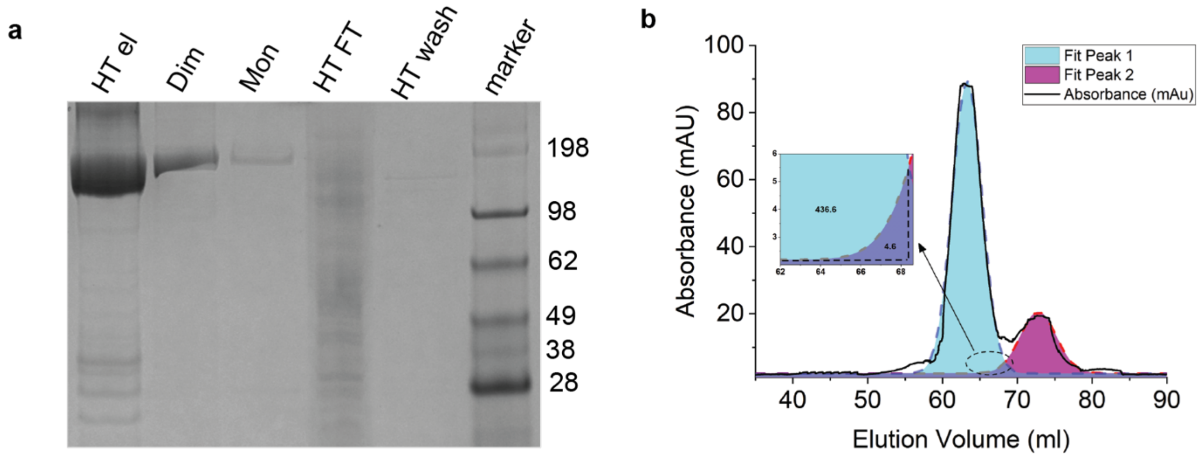


Supplementary figure 9: **α-catenin purification and dimer/monomer separation**. **a**, SDS-page gel, from the left: 1) elution of α-catenin from HisTrap columns; 2) dimeric and 3) monomeric fractions from size exclusion chromatography; 4) HisTrap flow-through; 5) HisTrap wash; 6) marker. **b**, Absorbance during elution from size exclusion column. The first peak from the left is from the dimeric α-catenin, the peak on the right from the monomeric α-catenin. From the area under the Gaussian fit of the two peaks (cyan and purple for the dimeric and monomeric catenin, respectively), we calculated the percentage of monomeric and dimeric catenin contained in each 1 ml elution aliquot. We pooled aliquots containing dimeric catenin with <1% monomeric catenin, and, similarly, monomeric catenin with <1% dimeric catenin. The figure inset shows the point where the monomeric catenin (area 4.6 AU) is about 1% of the dimeric catenin (area 436.6 AU).

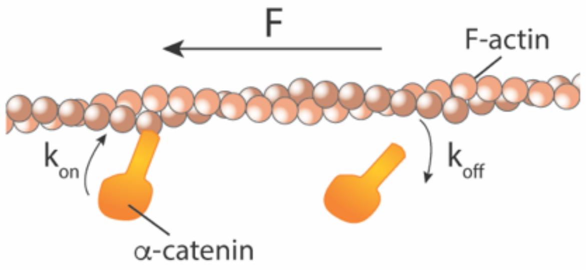


Supplementary figure 10: **α-catenin mechano-kinetic model.** Figure shows a sketch of the model of interaction of α-catenin with actin. The sliding velocity of the actin filament depends on the force applied to the filament (F) by active enzymes such as non-muscle myosin II and the force applied by α-catenin , which binds and unbinds from the filament with attachment and detachment rates $k_{on}$ and $k_{off}$, respectively

### Supplementary Methods

#### Pull down assay

A pull down assay was used to confirm the predicted interaction α-E-catenin and F-actin and measure the affinity of the complex *in vitro.* With such a method, proteins are mixed in solution and binding is evaluated in the absence of any external applied force (F=0). Each reaction contained increasing concentrations of α-catenin incubated with a fixed concentration of F-actin. Following binding and pelleting of the α-E-catenin-actin complexes through centrifugation, the proteins recovered from individual pull-downs were analyzed on an SDS-PAGE by densitometry (Supplementary Fig. 11a). Reactions and conditions are described in Methods. The intensities of the bands corresponding to the fraction of α-catenin bound to F-actin were quantified for each concentration. Data were well fitted by a Michaelis-Menten equation $I = \frac{I_{max}\cdot[\alpha cat]}{K_d\cdot[\alpha cat]}$, giving a dissociation constant $K_d$ = 0.65 ± 0.05 μM (Supplementary Fig. 11b), in good agreement with previous reports[1,2].

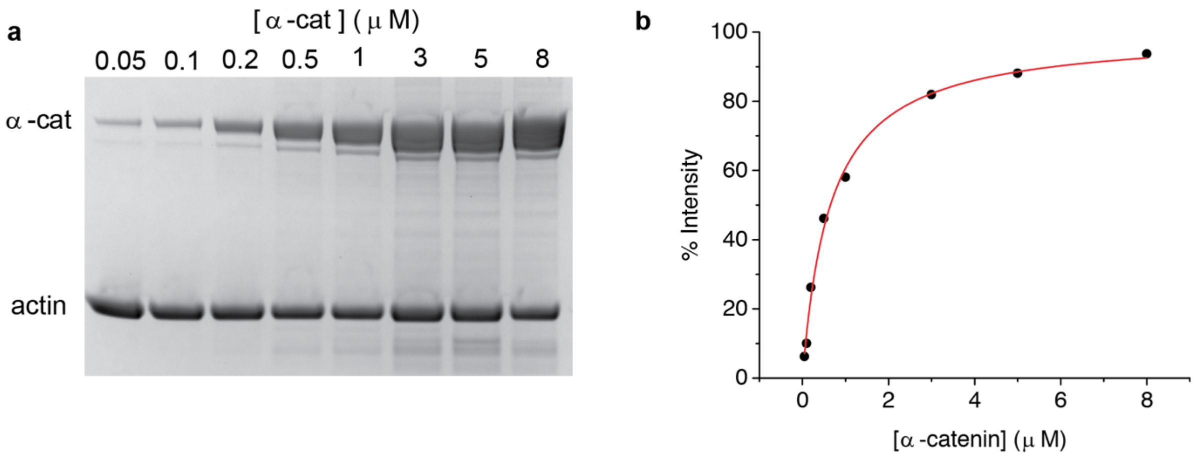


**Supplementary Fig. 11: α-catenin-actin pull-down assay. a,** SDS-page gel of the pellet of pull-down assay at increasing α-catenin concentration **b**, Densitometric analysis of the SDS-PAGE gel bands in (a) (black circles) was fitted with a Michaelis-Menten equation (red line).

#### Flow cell assay

Binding activity of α-catenin to F-actin was then evaluated through a flow cell assay to test whether α-catenin homodimers and α–β-catenin heterodimers could bind to actin under force. In this assay, α-catenin was first attached to the coverslip surface either on top of nitrocellulose or over a β-catenin bed. Next, fluorescently labelled F-actin was flowed and incubated into the chamber and then washed with an imaging buffer to remove floating F-actin and observe whether F-actin on the coverslip surface remained bound to α-catenin under the drag force applied by the buffer flow. The force applied to an actin filament by the buffer flow can be calculated as[3]:

$$\frac{F}{l} = c_{\parallel} v = \frac{2\pi\eta}{ln(2h/r)} v$$

where $l$ is the length of the actin filament, $c_{\parallel}$ is the drag coefficient per unit length along the actin filament axis, $v$ is the flow velocity (~2 mm/s), $\eta$ is the coefficient of viscosity of the buffer (~$10^{-3}$ N·s/m), $h$ is the distance between the coverslip surface and the center of the actin filament (~10 nm), and $r$ is the filament radius (~3 nm)[3] (Supplementary Fig. 12). From this formula we estimate a force per unit length of about 7 pN/μm. Given filament lengths in the range 1 - 10 μm, the flow cell assay allowed us to observe if multiple α-catenin molecules could bear forces of few tens of piconewton on actin.

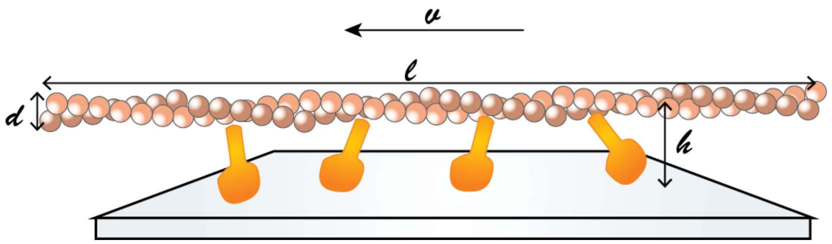


**Supplementary Fig. 12: α-catenin flow cell assay.** Figure shows a sketch of the experimental configuration of the flow cell assay indicating the distance between the coverslip surface and the center of the actin filament (h), the filament diameter (d), the length of the actin filament (l), and the buffer flow velocity.

Supplementary Fig. 13a and 13b show a field of view of a flow cell experiment in which α-catenin homodimers were bound onto a nitrocellulose smeared coverslip at concentration of 1 μM and 0.4 μM, respectively. Actin filaments bound on the surface were visible after the washing step in both conditions, with a significantly higher number of filaments for the higher α-catenin concentration. Supplementary Fig. 13c show a field of view of a flow cell experiment in which α-catenin at 1 μM concentration was bound onto a bed of β-catenin, attached to a nitrocellulose smeared coverslip. Also under these conditions, actin filaments bound on the surface were visible after the washing step. A control reaction in the absence of α-catenin was also performed showing no detectable F-actin filaments on the surface (Supplementary Fig. 13d). Reactions and conditions are described in detail in Methods.

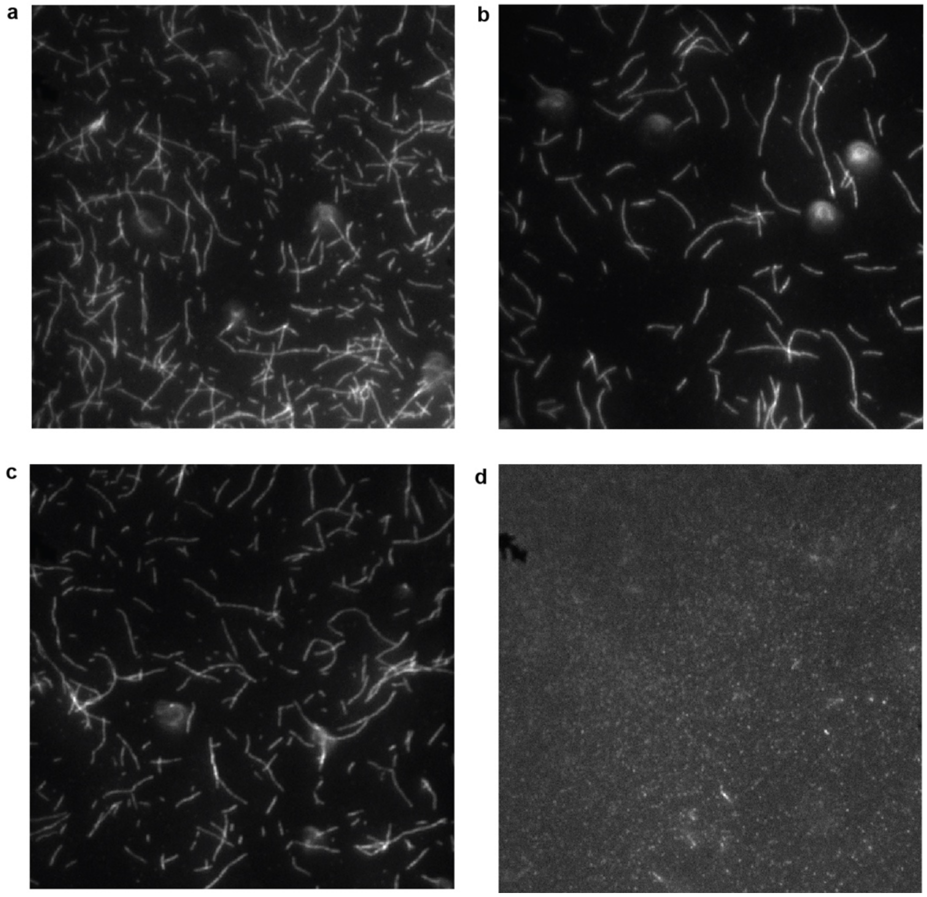


**Supplementary Fig. 13: α-catenin homodimer flow cell assay. a,** field of view of a flow cell experiment in which α-catenin homodimers at 1 μM concentration was bound onto a nitrocellulose smeared coverslip. Actin filaments bound on the surface were visible after the washing step. **b,** field of view of a flow cell experiment as in (a) but with α-catenin at 0.4 μM concentration **c,** field of view of a flow cell experiment in which α-catenin at 1 μM concentration was bound onto a bed of β-catenin, attached to a nitrocellulose smeared coverslip. **d**, Control reaction in the absence of α-catenin showing no detectable F-actin filaments on the surface.

#### Supplementary discussion

##### Step Size Distribution

In our experiments, a single α-catenin homodimer might bind to the actin filament with a single monomer or with both. Binding of both monomers to actin would lead to the possibility that the position steps observed at force > 5pN are the consequence of the unbinding of one of the two monomers. In fact, monomer unbinding would result in the displacement of the actin filament under force as a consequence of the change in the bond stiffness. Similarly, a step might result from the unbinding of one monomer in the experiments with multiple α-β-catenin heterodimers. However, we believe that such events cannot be the primary cause of the main peak observed in the step distributions, for the following reasoning.

Assume that the stiffness of a single monomer bound to F-actin is $k_m$ and the stiffness of the dimer bound to F-actin is $k_d = 2k_m$. Under a constant force $F$, the dimer would be strained by $x_d = F/k_d = F/2k_m$, whereas the monomer by $x_m = F/k_m$. Therefore, the step that we should observe when a monomer detaches from actin and the other stays bound is $d = x_m - x_d = F/2k_m$. Under this hypothesis, the $d$ ~ 12 nm step measured at $F$ ~ 5 pN force would imply that the stiffness of the monomer is about $k_m$ = 5 pN/(2*12 nm) = 0.21 pN/nm. At 11 pN force, the unbinding of one monomer would produce a step $d = F/2k_m$ = 11 pN / (0.42 pN/nm) = 26 nm, whereas we observe a main peak in the step distribution of similar size.

Therefore, in our opinion the most plausible explanation is that the step is due to a conformational change of the protein (unfolding) that does not modify substantially the protein stiffness. Under this hypothesis, since the applied force is constant and the protein is already strained by the force before the unfolding occurs (force application is much faster than the step dynamics[4]), the unfolding step would be independent of the applied force, as we observe.

It still remains the possibility that unbinding of one of the two monomers contributes to the larger steps that we see at larger forces. For example, at 11 pN force we observe a second peak around 35 nm, which might fit the sum of the unfolding step (12 nm) and monomer unbinding (26 nm).

##### Cooperative binding

It's well established that that the interaction between α-β-catenin heterodimers and actin is much weaker than between α-catenin homodimers and actin[5]. Different explanations have been proposed previously (i) the N- and C-terminal domains of α-catenin are allosterically coupled and binding to β-catenin on the N-terminal domain might alter the C-terminal domain ability to bind to actin[5]. (ii) Structural studies[6] indicate that β-catenin might sterically hinder F-actin binding by the α-catenin binding domain, which could be at the basis of the different F-actin binding between the homodimer and heterodimer. (iii) α-catenin ABD binding to actin is accompanied by a conformational change in the actin protomer that affects the filament structure. This alteration of the filament structure can be at the base of a cooperative binding mechanism that reinforces the link between an α-catenin homodimer and actin compared to an α-catenin monomer[7]. Our results indicate that a cooperative mechanism is at the basis of the bond reinforcement and the analysis of the α-catenin stiffness during the interaction with actin in the different experiments reinforces this interpretation (see discussion in the main text). However, the identification of the structural features that are at the basis of the different kinetics of α-catenin homodimers and heterodimers is out of the scope of our article, and further studies would be required to clearly assess this point.

**Non-specific interactions**

We made many control experiments to rule out non-specific interactions, which are one of the well-known issues in this kind of single molecule experiments. In experiments on α-β-catenin heterodimers, α-catenin was attached on the coverslip surface on top of a nitrocellulose-coated surface saturated with β-catenin, followed by BSA (see methods). We made several control slides in which the coverslip surface was coated as described above but in the absence of α-catenin and looked for non-specific interactions on several tens of beads in each slide. We could very rarely (less than one bead per slide) find non-specific interactions with this control surface. Moreover, non-specific interactions were very different from the interactions observed in the presence of α-catenin, showing few short interactions when the dumbbell was close to the coverslip surface and the actin filament was pushing on the bead (as detected from the change in the position signal) and disappeared when the dumbbell was moved slightly farther from the coverslip surface. On the other hand, in the presence of α-catenin at single molecule concentration, we observed interactions in one every 4 beads on average, the interactions were much longer at low forces (tens of milliseconds) and the number of interactions increased with force. A single molecule was able to produce as much as several tens of thousands interactions. The interactions were observed also when the actin filament was not pushing on the bead. This behavior was never observed in the absence of α-catenin.

**Dimeric vs monomeric α-catenin**

Before our experiments, we separated dimeric from monomeric α-catenin by using size exclusion chromatography. This procedure assures that less than 1% of the catenin was in a dimeric form in the experiments with α-β-catenin heterodimers (see supplementary Fig. 9). The concentration of the monomeric catenin that we used in the experiments at single molecule concentration was about 1 µg/ml (10 nM); in the ones at "high" concentration, catenin concentration was 10 µg/ml (100 nM). Since the dissociation constant of the α-catenin homodimer is 25 µM, at equilibrium about 0.04% and 0.4% would be dimeric at the single-molecule and high concentrations, respectively. Moreover, Pokutta et al. showed that the α-catenin homodimer does not bind to β-catenin even after overnight incubation[8]. Therefore, the few % contamination of dimeric catenin was most likely washed away after few minutes of incubation in the sample chamber (see methods).

#### Supplementary References


1. Rimm, D. L., Koslov, E. R., Kebriaei, P., Cianci, C. D. & Morrow, J. S. Alpha 1(E)-catenin is an actin-binding and -bundling protein mediating the attachment of F-actin to the membrane adhesion complex. *Proc. Natl. Acad. Sci.* (1995). doi:10.1073/pnas.92.19.8813
2. Hansen, S. D. *et al.* αE-catenin actin-binding domain alters actin filament conformation and regulates binding of nucleation and disassembly factors. *Mol. Biol. Cell* **24**, 3710–20 (2013).
3. Howard, J. *Mechanics of motor proteins and the cytoskeleton*. (Sinauer Associates, Inc. Publisher, 2001).
4. Capitanio, M. *et al.* Ultrafast force-clamp spectroscopy of single molecules reveals load dependence of myosin working stroke. *Nat. Methods* **9**, 1013–1019 (2012).
5. Drees, F., Pokutta, S., Yamada, S., Nelson, W. J. & Weis, W. I. α-Catenin Is a Molecular Switch that Binds E-Cadherin-β-Catenin and Regulates Actin-Filament Assembly. *Cell* **123**, 903–915 (2005).
6. Rangarajan, E. S. & Izard, T. Dimer asymmetry defines α-catenin interactions. *Nat. Struct. Mol. Biol.* **20**, 188–193 (2013).
7. Hansen, S. D. *et al.* αE-catenin actin-binding domain alters actin filament conformation and regulates binding of nucleation and disassembly factors. *Mol. Biol. Cell* **24**, 3710–20 (2013).
8. Pokutta, S., Choi, H.-J., Ahlsen, G., Hansen, S. D. & Weis, W. I. Structural and thermodynamic characterization of cadherin·β-catenin·α-catenin complex formation. *J. Biol. Chem.* **289**, 13589–601 (2014).
